# Integrated Multi-omic Framework of the Plant Response to Jasmonic Acid

**DOI:** 10.1101/736363

**Authors:** Mark Zander, Mathew G. Lewsey, Natalie M. Clark, Lingling Yin, Anna Bartlett, J. Paola Saldierna Guzmán, Elizabeth Hann, Amber E. Langford, Bruce Jow, Aaron Wise, Joseph R. Nery, Huaming Chen, Ziv Bar-Joseph, Justin W. Walley, Roberto Solano, Joseph R. Ecker

**Affiliations:** Plant Biology Laboratory, Salk Institute for Biological Studies, La Jolla, CA 92037, USA; Genomic Analysis Laboratory, Salk Institute for Biological Studies, La Jolla, CA 92037, USA; Howard Hughes Medical Institute, Salk Institute for Biological Studies, La Jolla, CA 92037, USA; Centre for AgriBioscience, Department of Animal, Plant and Soil Sciences, School of Life Sciences, La Trobe University, Melbourne, VIC 3086, Australia; Australian Research Council Industrial Transformation Research Hub for Medicinal Agriculture, Centre for AgriBioscience, La Trobe University, Bundoora, VIC 3086, Australia; Plant Pathology and Microbiology, Iowa State University, Ames, IA 50011, USA; School of Natural Sciences, University of California Merced, Merced, CA 95343, USA; Department of Chemical and Environmental Engineering, Department of Botany and Plant Sciences, University of California, Riverside, CA 92521, USA; Computational Biology Department, School of Computer Science, Carnegie Mellon University, Pittsburgh, PA 15213, USA; Department of Plant Molecular Genetics, Centro Nacional de Biotecnología, Consejo Superior de Investigaciones Científicas (CNB-CSIC), 28049 Madrid, Spain

## Abstract

Understanding the systems-level actions of transcriptional responses to hormones provides insight into how the genome is reprogrammed in response to environmental stimuli. Here, we investigate the signaling pathway of the hormone jasmonic acid (JA), which controls a plethora of critically important processes in plants and is orchestrated by the transcription factor MYC2 and its closest relatives in *Arabidopsis thaliana.* We generated an integrated framework of the response to JA that spans from the activity of master and secondary-regulatory transcription factors, through gene expression outputs and alternative splicing to protein abundance changes, protein phosphorylation and chromatin remodeling. We integrated time series transcriptome analysis with (phospho)proteomic data to reconstruct gene regulatory network models. These enable us to predict previously unknown points of crosstalk from JA to other signaling pathways and to identify new components of the JA regulatory mechanism, which we validated through targeted mutant analysis. These results provide a comprehensive understanding of how a plant hormone remodels cellular functions and plant behavior, the general principles of which provide a framework for analysis of cross-regulation between other hormone and stress signaling pathways.

## Introduction

Plant hormones are structurally unrelated small signaling molecules that play pivotal roles in a wide range of fundamental processes of plants spanning growth, development and responses to environmental stimuli (Vanstraelen and Benkova, 2012). Hormone perception by plants stimulates a cascade of transcriptional reprogramming that ultimately modifies cellular function and plant behavior (Chang et al., 2013, Song et al., 2016, Hickman et al., 2017, Pauwels et al., 2008). This is initiated by one or a family of high-affinity receptors, followed by signal transduction through protein-protein interactions, post-translational modification events and regulation of transcription factor (TF) activity that ultimately drives changes in gene expression (Wang et al., 2015, Song et al., 2016, Chang et al., 2013).

One of the key plant hormones is methyl jasmonate (JA), which regulates crucial processes including fertility, seedling emergence, the response to wounding and the growth-defense balance (Huang et al., 2017). Jasmonates are perceived as jasmonoyl-isoleucine (JA-Ile) by the co-receptor COI1 (CORONATINE INSENSITIVE1)/JAZ (Jasmonate ZIM domain) complex (Thines et al., 2007, Chini et al., 2007, Fonseca et al., 2009, Sheard et al., 2010). COI1 is an F-box protein and part of a Skp-Cullin-F-box-E3 ubiquitin ligase complex (SCF^COI1^) (Xie et al., 1998) that targets JAZ proteins for proteasomal degradation upon JA perception. JAZs are transcriptional repressor proteins that inhibit the activity of key TFs of the JA pathway such as the basic helix-loop-helix (bHLH) TF MYC2 and its closest homologues MYC3, MYC4 and MYC5 (Fernandez-Calvo et al., 2011, Song et al., 2017, Lorenzo et al., 2004) in the absence of JA. The SCF^COI1^-JAZ complex tightly controls the level of free non-repressed MYCs in a JA-dependent manner thereby determining the transcriptional output of the entire JA response (Chini et al., 2007, Thines et al., 2007, Zhang et al., 2015a). The key regulatory step in the JA pathway is the hormone-triggered formation of a complex between the E3 ligase SCF^COI1^ and JAZ repressors that are bound to the master TF MYC2. This results in degradation of JAZ repressors and permits the activity of a master regulatory bHLH transcription factor MYC2, accompanied by MYC3, MYC4, MYC5 and numerous other transcription factors, all of which have distinct but overlapping roles in driving JA-responsive gene expression (Song et al., 2017, Schweizer et al., 2013b, Fernandez-Calvo et al., 2011, Lorenzo et al., 2004, Bao et al., 2019). The result is a cascade of JA-induced genome reprogramming to modulate plant behavior such as plant immune responses (Du et al., 2017, Hickman et al., 2017). However, our knowledge of the JA-responsive genome regulatory program and, more broadly, in the plants general response to environmental stimuli is limited currently by assessment of only one or a small number of components.

Here we aim to decipher MYC2/MYC3-driven JA-responsive gene expression using a multi-omics analysis that includes the direct targets of key transcription factors, chromatin modifications, global protein abundance and protein phosphorylation. We discovered that MYC2/MYC3 directly target hundreds of TFs, resulting in a large transcriptional network that facilitates extensive crosstalk with other signaling pathways. This model predicted new components of the JA signaling pathway that we validated by targeted genetic analyses, demonstrating the power of our integrated multi-omic approach to yield fundamental biological insight into plant hormone responses.

## Results

### MYC2 and MYC3 binding is associated with a large proportion of JA-responsive genes

To decipher the JA-governed regulatory network with its high degree of dynamic and spatio-temporal interconnectivity with other signaling pathways, we applied a multi-omic network approach that is comprised of five newly generated large-scale datasets (Fig. 1a, b). MYC2 is the master regulatory transcription factor of JA responses and plants with a null mutation causing a clear decrease in JA sensitivity (Lorenzo et al., 2004). Thus, we included the *myc2* (*jin1-8* SALK_061267) mutant into our analyses (Fig. 1b) (Lorenzo et al., 2004). MYC2 is responsible for strong JA-responsive gene activation and acts additively with MYC3 and MYC4 (Lorenzo et al., 2004, Fernandez-Calvo et al., 2011). *myc3* and *myc4* single mutants behave like wildtype with regards to JA-induced root growth inhibition. However, in combination with the *myc2* mutant, *myc2 myc3* double mutants exhibit an increased JA hyposensitivity, almost as pronounced as in *myc2 myc3 myc4* triple mutants (Fernandez-Calvo et al., 2011). We consequently selected MYC3 for an in-depth analysis.

**Figure 1.**
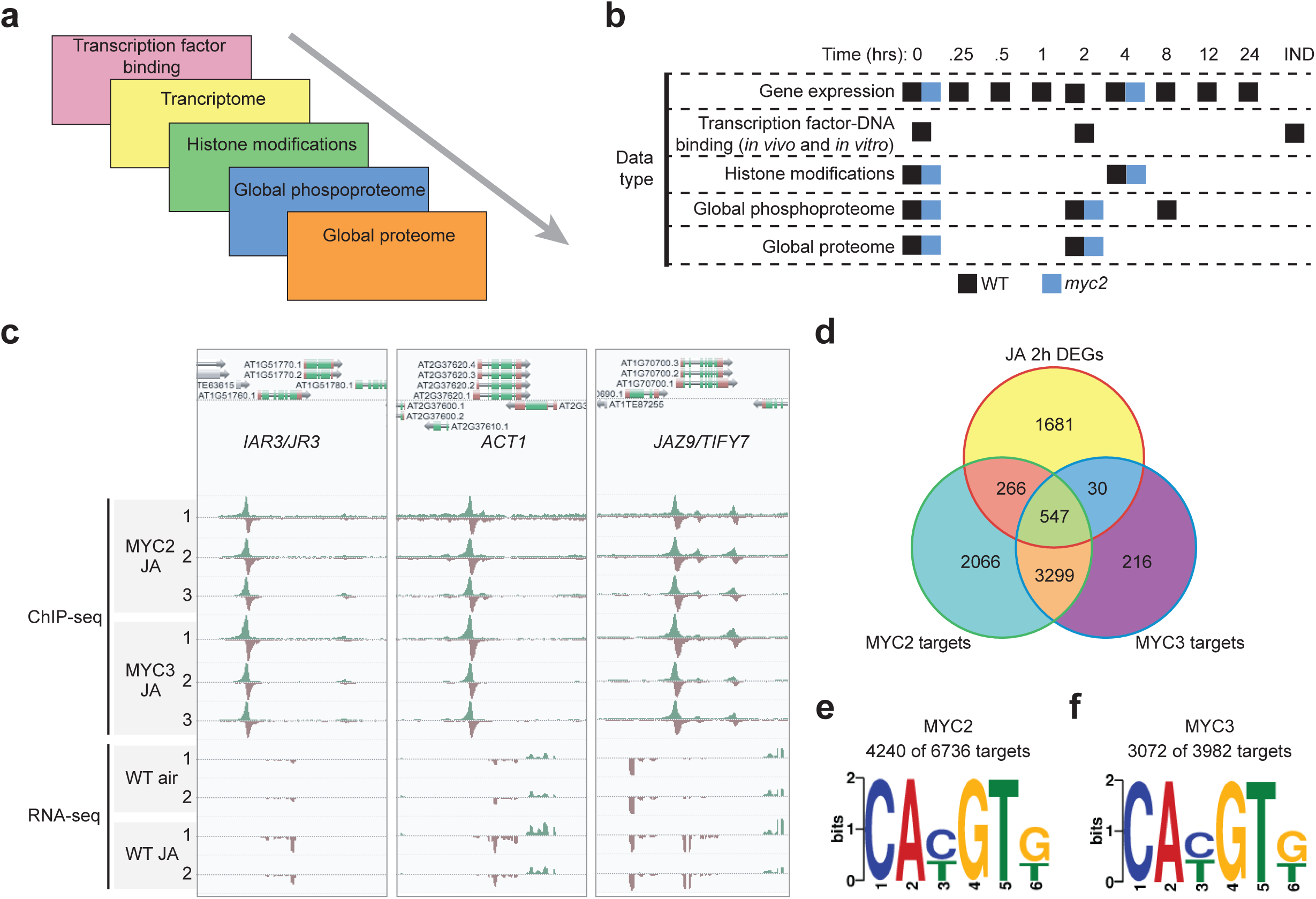
Design of our study and key datasets utilized. **a, b,** Overview of profiled regulatory layers (a) and detailed description of collected samples (b). **c,** AnnoJ genome browser screenshot visualizes the binding of MYC2 and MYC3 to the three example genes (I*AR3/UR3*, *ACT1*, *JAZ9/TIFY7*). MYC2/3 binding was determined with ChIP-seq using JA-treated (2 hours) Col-0 *MYC2::MYC2-Ypet* and Col-0 *MYC3::MYC3-Ypet* seedlings. Three independent biological ChIP-seq replicates are shown. In addition, mRNA expression of the three example genes Col-0 seedlings (-/+ 2 hours JA) is shown as well. Expression data is derived from RNA-seq analysis. **d,** Venn diagram illustrates the overlap between MYC2, MYC3 target genes and genes that are differentially expressed after two hours of JA treatment (JA 2h DEGs). **e, f,** The top-ranked motif in MYC2 (e) and MYC3 (f) ChIP-seq data was the G-box (CAC/TGTG. Motifs were determined by MEME analysis using the top-ranked peaks that were identified by GEM Motifs enriched in MYC2 and MYC3 peaks.

In order to better understand how the master TFs MYC2 and MYC3 control the JA-induced transcriptional cascade, we determined their genome-wide binding sites using chromatin immunoprecipitation sequencing (ChIP-seq). Four biological replicates of JA-treated (2 hours) three-day-old etiolated *Arabidopsis* seedlings that express a native promoter-driven and epitope (YPet)-tagged version of MYC2 and three biological replicates of MYC3 (Col-0 *MYC2::MYC2-Ypet*, Col-0 *MYC3::MYC3-Ypet*) were used (Gimenez-Ibanez et al., 2017). The genome-wide distributions of MYC2 and MYC3 binding sites were highly similar (Fig. 1c, d). We identified 6,736 MYC2 and 3,982 MYC3 binding sites of high confidence, equating to 6,178 MYC2 and 4,092 MYC3 target genes (Fig. 1d and Supplementary Table 1). Of the target genes identified, 3,847 were shared, meaning that almost all MYC3 target genes are also bound by MYC2 (Fig. 1d). Their target genes were enriched for JA-related gene ontology terms and for terms related to other hormones (Zhang et al., 2014, Abe et al., 2003) (Supplementary Fig. 1a). Collectively, these data indicate that MYC2 and MYC3 have the potential to regulate 23.2% of genes in the *Arabidopsis* genome (27,655 coding genes). However, binding events are not necessarily regulatory (Chang et al., 2013, Song et al., 2016, Fernandez et al., 2003). We determined that 2,522 genes are differently expressed (false discovery rate, FDR <0.05) after two hours of JA treatment using RNA-seq. A third (843 genes) of JA-modulated genes were directly bound by MYC2 or MYC3 (Fig. 1d and Supplementary Table 2). This is consistent with the important role of MYC2/3 in JA-responsive gene expression (Lorenzo et al., 2004, Fernandez-Calvo et al., 2011, Schweizer et al., 2013b). The majority of JA-responsive direct MYC2/3 target genes are transcriptionally upregulated after JA application indicating that MYC2 and MYC3 predominantly act as transcriptional activators (Supplementary Fig. 1b).

The G-Box (CAC/TGTG) was the most common DNA sequence motif found at MYC2 or MYC3 binding sites, which is concordant with the observation that they shared a large proportion of their binding sites (Fig. 1e, f). This motif was also of similar sequence to a motif bound by MYC2 determined *in vitro* (Godoy et al., 2011). The majority of MYC2 and MYC3 binding sites contained the G-Box motif (MYC2: 4,240 of 6,736; MYC3: 3,072 of 3,982; Fig. 1e, f and Supplementary Table 3). However, the absence of the motif from a substantial number of MYC2 and MYC3 binding sites suggests the transcription factors may bind indirectly to some sites through partner protein(s).

Master TFs directly target the majority of signaling components in their respective pathway, a phenomenon which has already been observed already for the ethylene, abscisic acid and cytokinin signaling pathways (Chang et al., 2013, Song et al., 2016, Xie et al., 2018). This pattern also holds true for the JA signaling pathway. Our MYC2/MYC3 ChIP-seq analyses determined that approximately two thirds of genes encoding for known JA pathway components (112 of 168 genes for MYC2 and 96 of 168 genes for MYC3) are bound by MYC2 and MYC3 (Supplementary Fig. 1c, d and Supplementary Table 4). Interestingly, the majority of all known JA genes that were differentially expressed following JA treatment were bound by MYC2 or MYC3 whereas fewer non-differentially expressed known JA genes were directly targeted (Supplementary Fig. 1c and Supplementary Table 4). MYCs initiate various feed forward loops that allow a rapid activation of the transcriptional JA response (Du et al., 2017, Liu et al., 2019). Our ChIP-seq approach revealed that besides the autoregulation of MYC2 and MYC3, they also regulate JA biosynthesis either indirectly through binding to the AP2-ERF transcription factor gene *ORA47* (Chen et al., 2016a) or directly by targeting the JA biosynthesis genes *LOX2* and *AOS2* (Supplementary Table 4). In addition, MYCs simultaneously target various negative regulators enabling MYCs to efficiently dampen the JA response pattern. Key negative regulators of JA signaling are the JAZ repressors, a gene family of 13 members in *Arabidopsi*s (Guo et al., 2018, Chung et al., 2010, Cuellar Perez et al., 2014) which can interact with the adaptor protein NINJA to confer TOPLESS-mediated gene repression (Pauwels et al., 2010). Strikingly, all JAZs and also NINJA are directly bound by MYC2 and MYC3 (Supplementary Fig. 1e), with the probable effect of dampening the JA response thereby preventing excessive activation of JA signaling.

### MYC2 and MYC3 activate the transcriptional JA response through a large transcription factor network

To decipher the MYC2 and MYC3-governed transcriptional regulatory network in more detail, we investigated the relationship between MYC2/MYC3-bound TF-encoding genes and their transcriptional responsiveness to JA treatment. We conducted a JA time course experiment (time points 0, 0.25, 0.5, 1, 2, 4, 8, 12, 24 h post JA treatment) identifying a total of 7,377 differentially expressed genes at one or more time points within 24 h of JA treatment (Supplementary Table 2). Differentially expressed genes were categorized into clusters with similar expression trends over time to facilitate visualization of complex expression dynamics and enriched functional annotations (Supplementary Fig. 2a and Supplementary Table 5). The largest upregulated cluster was the “JA cluster” which was enriched for gene ontology (GO) terms associated with JA responses (Fig. 2a). In contrast, the “Cell wall cluster” was the largest cluster of downregulated genes and enriched for GO terms associated with cell wall organization, development and differentiation (Fig. 2b). These two main clusters illustrate the defense-growth trade-off that plants are faced when defense pathways are activated (Huot et al., 2014).

**Figure 2.**
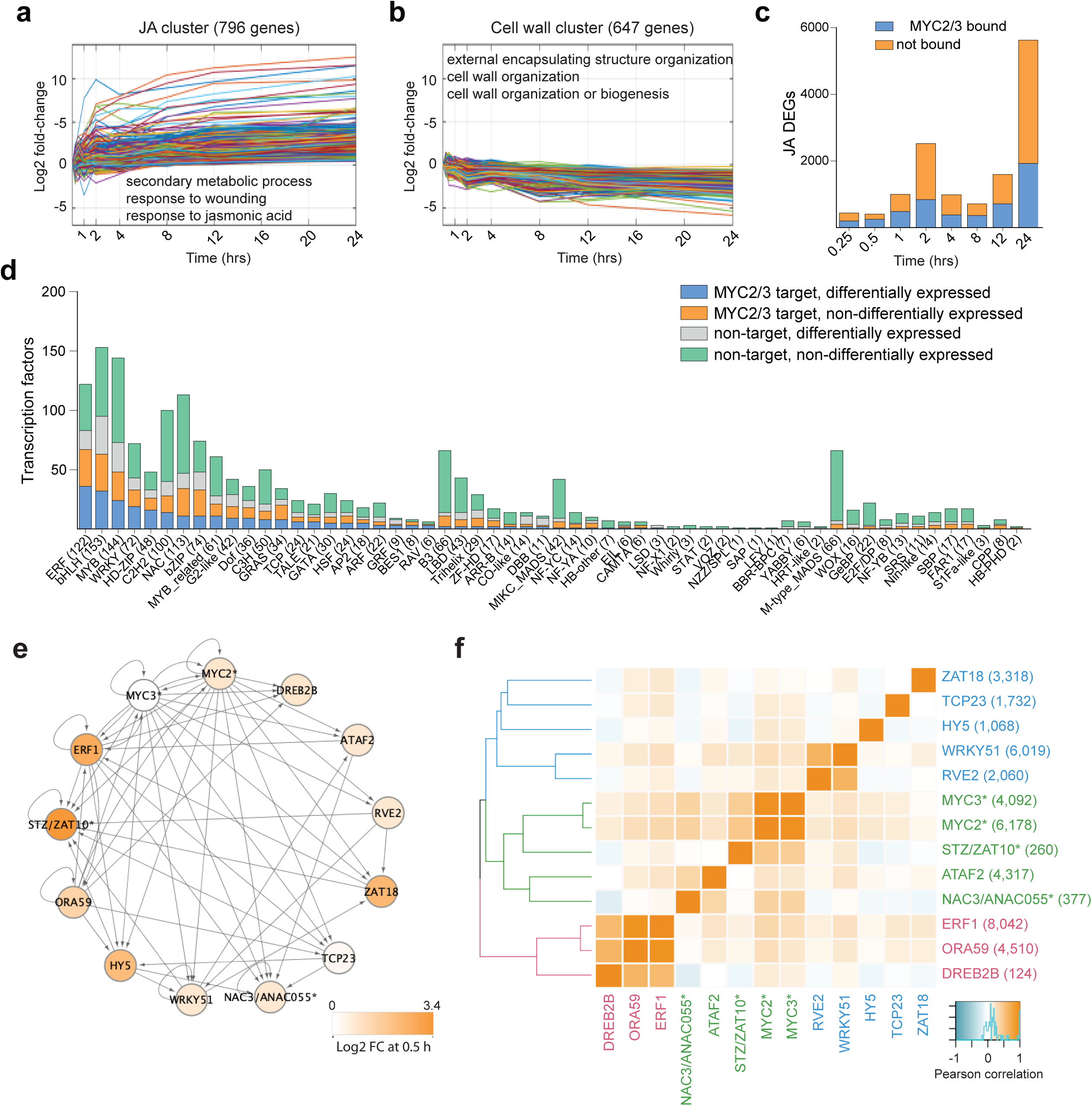
MYC2 and MYC3 target a large proportion of JA-responsive genes that encode transcription factors. **a, b,** Cluster analysis revealed the two main clusters in the JA time course experiment. The JA cluster (a) with 796 genes reflects the majority of JA-induced genes and the cell wall cluster (b) with 647 genes represents the largest cluster of JA repressed genes. Clusters visualize the log2 fold change expression dynamics over the indicated 24 hours’ time period. The three strongest enriched gene ontology terms for each cluster are shown as well. **c,** Bar plots illustrates the portion of JA differentially expressed genes (JA DEGs) that are bound by MYC2 and/or MYC3 at the indicated time points. JA DEGs for all time points were identified with RNA-seq. MYC2/3 targets are derived from ChIP-seq analysis using Col-0 *MYC2::MYC2-Ypet* and Col-0 *MYC3::MYC3-Ypet* seedlings that were treated for two hours with JA. **d,** MYC2 and MYC3 target genes from a wide range of transcription factor TF families. TF families are classified into four different groups; MYC2/MYC3 targets and differentially expressed after JA treatment (blue), MYC2/MYC3 targets and not differentially expressed (orange), not bound by MYC2/MYC3 but differentially expressed (grey) and not bound by MYC2/MYC3 but not differentially expressed (green). **e,** Nodes represent JA TFs for which direct binding data was generated. ChIP-seq data is indicated by presence of *, all other data was DAP-seq. Edges represent binding events and are directed. Self-loops indicate TF binds to its own locus, indicative of potential auto-regulation. Expression of the TF at 0.5 h after JA treatment is represented by color scale. **f,** Pearson correlation of TFs’ target sets of genes. Numerals in brackets indicate total number of target genes

Up to 63% (0.5 h JA treatment) of differentially expressed genes at any given time point were directly bound by MYC2 and/or MYC3 (Fig. 2c), highlighting the important role of MYCs in transcriptionally regulating JA responses. Our analysis also determined that 522 of 1,717 known or predicted TFs were differentially expressed within 24 h of JA treatment (Supplementary Fig. 2b). Half of these (268), representing 36 of 58 TF families, were also direct MYC2 or MYC3 targets (Fig. 2d and Supplementary Fig. 2b) indicating that MYC2 and MYC3 cooperatively control a massive TF network. The three most numerous families (ERFs, bHLHs and MYBs) in the *Arabidopsis* genome had the most JA-responsive MYC2 or MYC3 targeted members which is concordant with their previously annotated roles in JA responses (Fig. 2d) (Chen et al., 2016b). Plant hormone crosstalk is critical for an appropriate cellular response to environmental stimuli and numerous reports describe that MYC2 connects the JA pathway to other major plant hormone pathways (Hou et al., 2010, Lorenzo et al., 2004, Aleman et al., 2016, Zhang et al., 2014, Cui et al., 2018, Pieterse et al., 2009). To investigate this crosstalk function of MYC2 and MYC3 in more detail, we utilized our ChIP-seq data to determine the number of plant hormone TFs that are bound by MYC2 and MYC3. We found that 37 to 59% of annotated hormone pathway genes are bound by MYC2 and MYC3 and that their expression changes in response to 24 hours of JA treatment (Supplementary Fig. 2c). In addition, we discovered 122 annotated hormone TFs, with representatives from all hormone pathways, that are bound by MYC2 and MYC3 and 118 of these are differentially expressed (Supplementary Fig. 2d and Supplementary Table 1).

We next set out to better understand the target genes of the network of TFs downstream of MYC2 and MYC3. To do so we conducted ChIP-seq or DNA affinity purification sequencing (DAP-seq) (O’malley et al., 2016, Bartlett et al., 2017), on a subset of TFs that were direct MYC2/3 targets and rapidly upregulated (within 0.5 h) by JA treatment (DREB2B, ATAF2, HY5, RVE2, ZAT18; Fig. 2e) or were members of the upregulated “JA cluster” (TCP23; Fig. 2a). We also included TFs with known roles in JA signaling (ERF1, ORA59, NAC3/ANAC055, WRKY51, ZAT10) (Lorenzo et al., 2003, Pre et al., 2008, Bu et al., 2008, Gao et al., 2011, Pauwels and Goossens, 2008). These TFs formed a highly connected network, with all TFs except DREB2B targeting at least two TFs in the network and being themselves targeted by two TFs (Fig. 2e and Supplementary Table 6). Auto-regulation was common, with seven TFs targeting their own loci (Fig. 2e). The target genes of ZAT10, ANAC055 and ATAF2 were most similar to those of MYC2/3 (Fig. 2f). Consistent with this, their target genes shared several significantly enriched gene ontology terms (adjusted p<0.05), suggesting related functions in jasmonate signaling (Supplementary Fig. 2d). ORA59 and ERF1, along with DREB2B, formed a distinct group that targeted a related set of genes (Supplementary Fig. 3a). Notably, ERF1 and ORA59 also shared significant enrichment of a separate set of gene ontology terms with one-another, but that were not enriched amongst MYC2/3 targets. This is consistent with the joint role of ERF1 and ORA59 in controlling a pathogen defense arm of JA signaling (Pre et al., 2008, Lorenzo et al., 2003). No gene ontology terms were enriched amongst the targets of DREB2B. WRKY51 and RVE2 had relatively few enriched gene ontology terms but shared most of these with one-another. Most of the terms related to anti-insect defense and were a subset of the enriched MYC2/3-ZAT10-ANAC055-ATAF2 gene ontology terms (Supplementary Fig. 2a). ZAT10 and ANAC055 are known regulators of anti-insect defense and our results suggest WRKY51 and RVE2 may also be involved in this component of jasmonate responses (Schweizer et al., 2013a). Taken together, our analyses determine that MYC2 and MYC3 shape the dynamic spatiotemporal JA response through the activation of a large TF network that includes various potentially coupled feedforward and feedback loops and that allows extensive cross-communication with other signaling pathways.

### MYC2 controls JA-induced epigenomic reprogramming

Reprogramming of the epigenome is an integral part of development and environmental stimulus-induced gene expression (Feng et al., 2010, Xiao et al., 2017). For example, activation of the transcriptional JA response requires the formation of MYC2/MED25-mediated chromatin looping (Wang et al., 2019). To investigate the extent of JA-induced changes in chromatin architecture and the regulatory importance of MYC2 in this response, we conducted ChIP-seq assays to profile the genome wide occupancy of the histone modification H3K4me3 and the histone variant H2A.Z in untreated/JA-treated (4 h) Col-0 and *myc2* seedlings. Trimethylation of H3K4me3 marks active and poised genes and the histone variant H2A.Z confers gene responsiveness to environmental stimuli (Rothbart and Strahl, 2014, Coleman-Derr and Zilberman, 2012). mRNA expression was monitored in parallel using RNA-seq. JA treatment led to a reprogrammed chromatin landscape with several thousand differentially enriched H3K4me3 and H2A.Z domains (Supplementary Fig. 4a, b, c and Supplementary Table 7). We identified 826 differentially expressed genes (675 induced, 151 repressed; Col-0 control *v.* JA-treated) in that experiment and, as expected, the JA-induced genes had a stronger promoter enrichment of MYC2 than the JA-repressed genes (Fig. 3a and Supplementary Table 2). The JA-induced genes had an increase of H3K4me3, whereas JA-repressed genes had no dynamic change in the level of H3K4me3 (Fig. 3b, d). Strikingly, *myc2* mutants only display a compromised increase of H3K4me3 after JA treatment suggesting that the JA-induced trimethylation of H3K4me3 strongly depends on prior MYC2 binding (Fig. 3b, c, d and Supplementary Fig. 4a). This scenario is illustrated by two JA-induced genes, *JAZ2* and *GRX480*, which are directly targeted by MYC2. Their expression depends on MYC2 and their JA-induced increase of gene body-localized H3K4me3 partially depends on MYC2 (Fig. 3d and Supplementary Fig. 4d). In contrast, JA-induced changes in H2A.Z occupancy are not affected in *myc2* mutants (Supplementary Fig. 4a) suggesting that JA-induced H2A.Z dynamics are either independent of MYC2 or other MYCs such as MYC3, MYC4 and MYC5 are functionally redundant in regulating H2A.Z dynamics.

**Figure 3.**
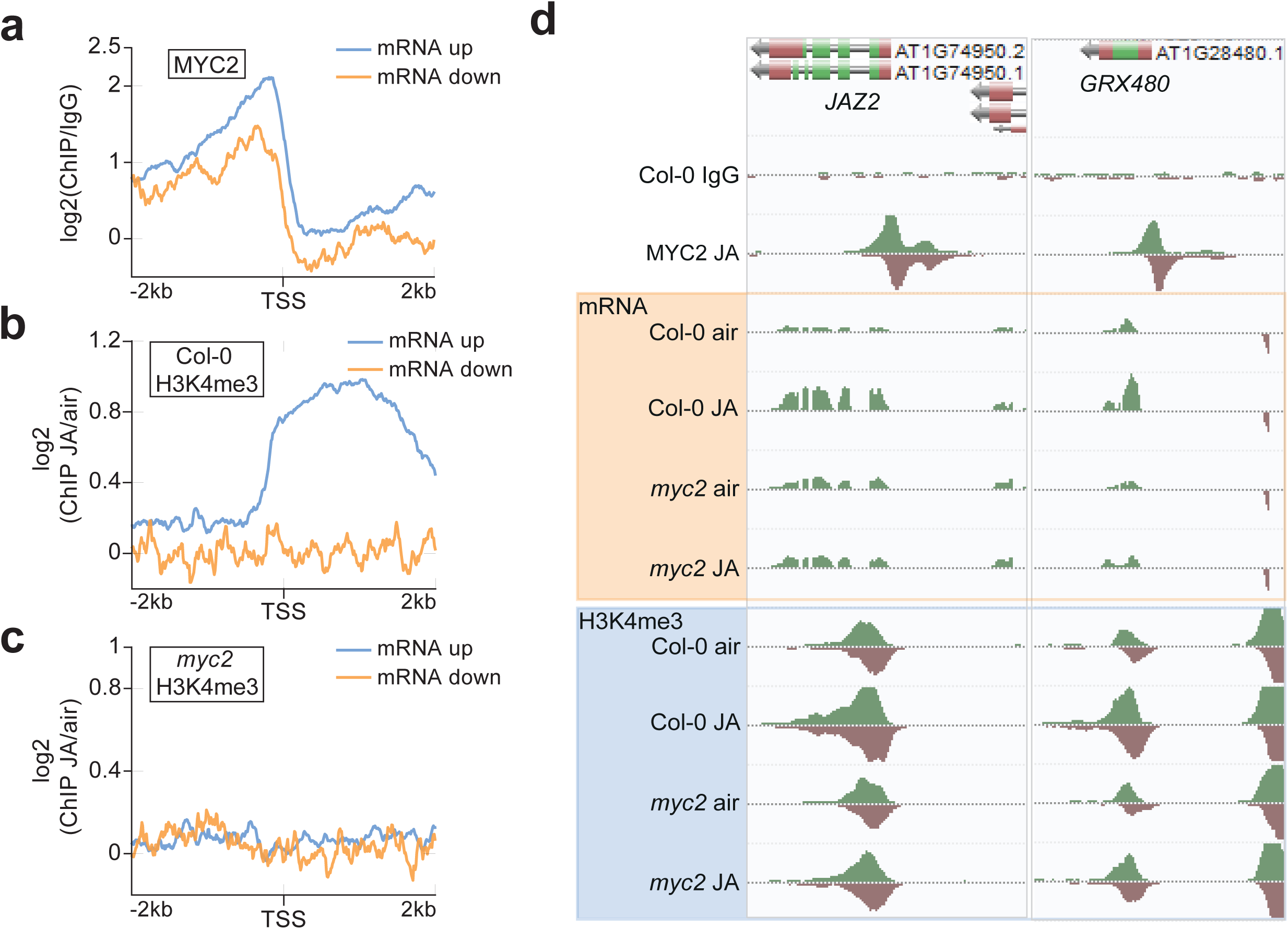
The jasmonic acid-responsive epigenome. **a, b, c,** Aggregated profiles show the log_2_ fold change enrichment of MYC2 (a) and H3K4me3 (b, c) from 2 kb upstream to 2 kb downstream of the transcriptional start site (TSS) at JA-induced and JA-repressed genes. Profile of MYC2 is shown for Col-0 *MYC2::MYC2-Ypet* (a) seedlings and H3K4me3 profiles are shown are shown for Col-0 (b) and *myc2* (c) seedlings. **d,** AnnoJ genome browser screenshot visualizes MYC2 binding, mRNA expression and H3K4me3 occupancy at two example genes (*JAZ2, GRX480*) in Col-0 and *myc2* seedlings. All tracks were normalized to the respective sequencing depth.

### Extensive remodelling of the (phospho)proteome occurs following a JA stimulus and may drive alternative splicing

We next explored how JA remodels the proteome and phosphoproteome of etiolated seedlings. Hormone signal transduction typically modifies phosphorylation of downstream proteins, changing their activity independent of transcript abundance (Wang et al., 2015). Transcript abundances are also frequently weakly correlated with protein abundances (Walley et al., 2016, Baerenfaller et al., 2008). Consequently, proteomic and phosphoproteomic analyses yield additional insight into gene regulatory networks. We determined that the loss of MYC2 caused substantial changes to the JA-responsive proteome and phosphoproteome; 1,432 proteins and 939 phosphopeptides (corresponding to 567 genes) were significantly differentially abundant in Col-0 seedlings relative to *myc2* seedlings after 2 h JA treatment (q<0.01; Fig. 4a and Supplementary Table 8, 9). Col-0 seedlings responded to JA (161 proteins, 443 phosphopeptides, Col-0 JA v. Col-0 air) and the response was smaller without functional MYC2 (79 proteins, 93 phosphopeptides, *myc2* JA v. *myc2* air). Some overlap existed between proteins or phosphopeptides and transcripts responsive to JA treatment (Fig. 4b). Both transcripts and proteins encoded by 28 genes were differentially expressed in JA-treated Col-0 seedlings relative to air controls. A further 33 differentially expressed proteins in JA-treated Col-0 seedlings had no corresponding differentially expressed transcript but were encoded by genes that are targeted by MYC2 and MYC3. Differentially abundant phosphopeptides were detected that corresponded with 15 differentially expressed transcripts. Transcript and protein abundances also correlated poorly in JA-treated Col-0 seedlings (Fig. 4c), in agreement with prior studies (Walley et al., 2016, Baerenfaller et al., 2008). The protein of only one known JA pathway component was differentially abundant in JA-treated Col-0 seedlings relative to controls, and none were differentially phosphorylated. In sum, these data indicate that the JA-responsive proteome and phosphoproteome are poorly annotated and are not well represented by transcriptome studies (Fig. 4c).

**Figure 4.**
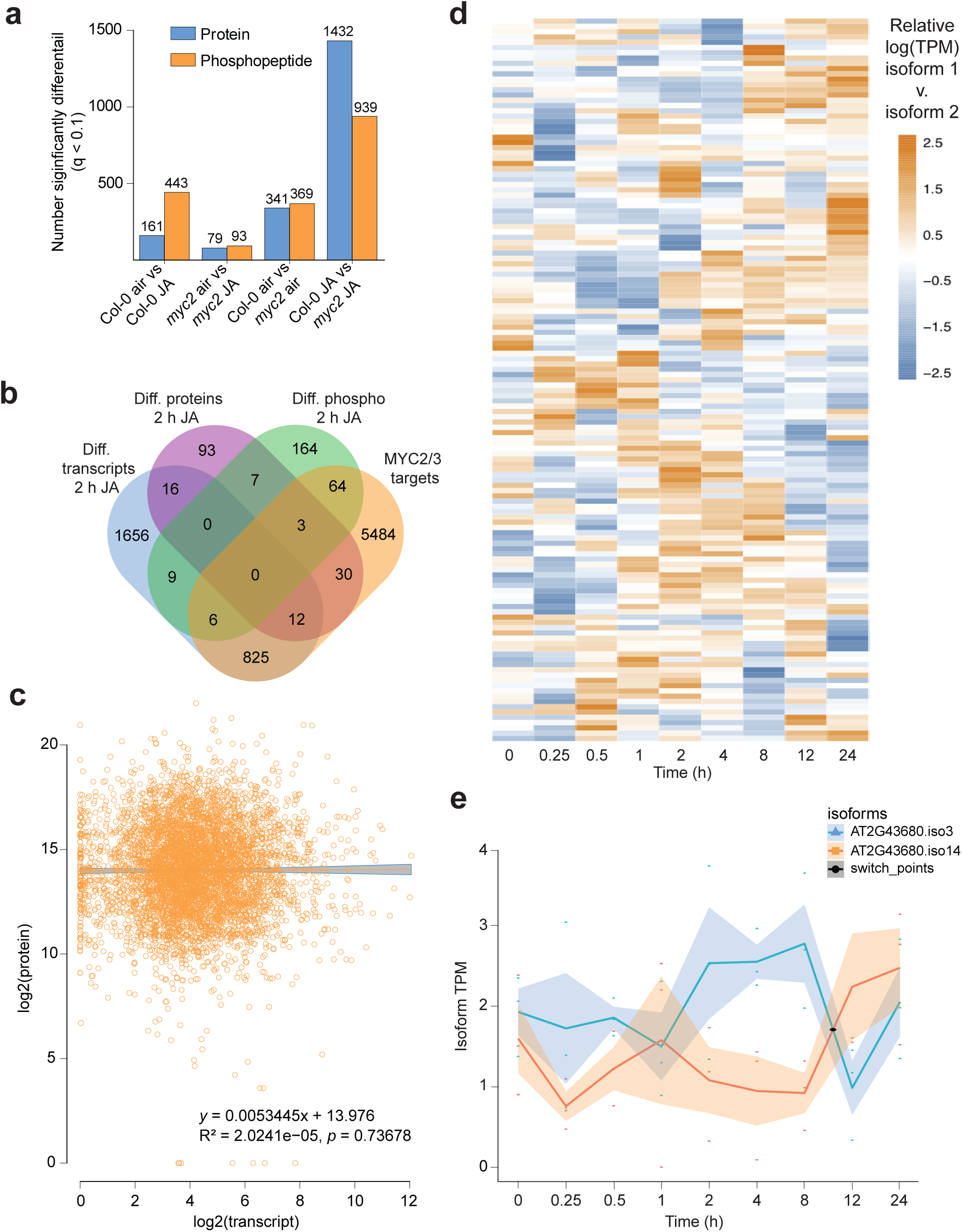
Loss of functional MYC2 impacts the global proteome and phosphoproteome. **a,** Total significantly differentially abundant (q<0.1) proteins and phosphopeptides detected in comparisons between JA-treated (2 h) Col-0 and *myc2* seedlings and mock controls. **b,** Venn diagram showing the overlap between significantly differentially abundant proteins, transcripts and differentially phosphorylated proteins in JA-treated Col-0 seedlings compared to mock-treated Col-0 controls. Also shown is the overlap with MYC2/3 target genes. **c,** Correlation between log_2_(FPKM)s of detected proteins and transcripts in Col-0 seedlings treated with JA for 2 h. Scatter plot of log_2_ fold change in Col-0 JA-regulated transcript levels versus log_2_ fold change in levels of corresponding proteins. **d,** Heatmap represents relative TPM of 137 isoform pairs exhibiting isoform switch events. Ratio calculated as logTPM (isoform 1/isoform 2). **e,** Plot shows an example of a transcript pair originating from *AT2G43680* that had isoform switch events following JA treatment.

Alternative splicing can occur rapidly in response to environmental stimuli, contributing to transcriptome reprogramming and potentially fine-tuning physiological responses (Hartmann et al., 2016, Calixto et al., 2018). It is central to JA-mediated regulation of transcription, with an alternative isoform of the repressor JAZ10 creating a negative feedback loop that desensitizes cells to a JA stimulus (Moreno et al., 2013, Zhang et al., 2017, Chung et al., 2010). However, the extent of alternative splicing in JA signaling beyond the JAZ repressors is poorly characterized. We observed that phosphorylation of proteins involved in RNA recognition and nucleotide binding was disrupted in JA-treated *myc2* mutants compared with Col-0 seedlings. The spliceosome was the only pathway significantly enriched amongst these differentially phosphorylated proteins (p < 0.05, 18 genes matched) suggesting that MYC2 may influence JA-responsive alternative splicing. We examined isoform switching events across our JA transcriptome time-series, where the most abundant of two isoforms from a single gene changes, to determine the extent of JA-responsive alternative splicing (Fig. 4d, e, Supplementary Table 10). There were 151 switch events, corresponding to 137 isoform pairs from 120 genes, within 24 h of JA treatment. These were identified from 30,547 total individual transcripts detected (average TPM>1; Supplementary Table 11). Two of the genes exhibiting isoform switches had prior JA annotations (*RVE8/AT3G09600*, *SEN1/AT4G35770*) and others were annotated to a variety of processes (including auxin, ABA, light signaling, disease response, amongst many others), but there was no significant enrichment of any gene ontology terms or pathways. This indicates that MYC2 influences alternative splicing that diversifies the transcriptome in response to a JA stimulus.

### Multi-omic modelling provides a comprehensive understanding of the JA response genome regulatory program

We wanted next to characterize the broader JA-response genome regulatory program so that we could increase our understanding of the roles of known JA TFs within this and identify new potential regulatory interactions. To do so we generated a gene regulatory network model encompassing the (phospho)proteomic and time-series transcriptomic data (Supplementary Fig. 5a and Supplementary Table 12). Many known JA signaling components were present in the 100 most important predicted components of the model (MYC2, ERF1, JAZ1, JAZ2, JAZ5, JAZ10, ATAF2 and others; within top 100 of 4366 components by normalized motif score; Supplementary Table 12). MYC2 was predicted to regulate a subnetwork of 26 components, 23 of which were validated as directly bound by MYC2 in ChIP-seq assays (88.5%, Supplementary Fig. 6a and Supplementary Table 1, 12). We further validated the network by comparing the ChIP/DAP-seq data previously collected for the remaining 12 JA TFs to their targets in the gene regulatory network (Fig. 2, Supplementary Fig. 6b and Supplementary Table 13). The gene regulatory network identified all of these TFs as components of the JA response, except MYC3 (Supplementary Table 12). It is likely that MYC3 was not part of the network due to it being only modestly differentially expressed following JA treatment and not being detected in the (phospho)proteome analyses (Supplementary Tables 2, 8, 9). The wider validation of targets was less strong than for MYC2, ranging from 0% to 33.3%. This could reflect the possibility that interactions predicted by the gene regulatory network may not identify all intermediate components. Lastly, we examined known genetic interactions. The MYC2 subnetwork included activation of JAZ10 within 0.5 h of a JA stimulus, with JAZ10 reciprocally repressing MYC2 (Supplementary Fig. 6a, b). This is consistent with the known role of JAZ10 in establishing negative feedback that attenuates JA signaling (Moreno et al., 2013). MYC2 was also predicted to activate AIB (JAM1/bHLH017/AT2G46510) (Supplementary Fig. 6a, b), establishing a negative feedback loop in which AIB negatively regulates MYC2. This is consistent with prior studies, which established AIB is dependent upon and antagonistic to MYC2, thereby repressing JA signaling (Nakata et al., 2013, Sasaki-Sekimoto et al., 2013, Fonseca et al., 2014). Confirmation by both genetic data from the literature and our DAP/ChIP-seq experiments indicates that our gene regulatory network modelling approach is a useful tool to identify new regulatory interactions within JA signaling and to better understand known regulatory interactions.

Crosstalk between hormone response pathways permits fine-tuning of plant growth and development in response to diverse environmental signals (Karasov et al., 2017). We examined the potential points at which MYC2 may interface directly with other hormone signaling pathways, since MYC2 is the master regulator of JA responses and one of the first TFs activated by JA. The MYC2 subnetwork identified a potential route for JA signaling to cross-regulate auxin hormone signaling. MYC2 activated ARF18 and ARF18 reciprocally activated MYC2 (Supplementary Fig. 6a and Supplementary Table 12). It also indicated that MYC2 may promote ethylene signaling by activating MAP kinase kinase 9 (MKK9) (Supplementary Fig. 6a). Prior genetic studies determined that MKK9 induces ethylene production, but had not examined a possible link with JA signaling (Xu et al., 2008). Positive crosstalk is known to exist between JA and auxin signaling though the mechanism is not clearly determined (An et al., 2010, Hentrich et al., 2013). RGL3, a regulator of gibberellic acid (GA) signaling previously associated with JA-GA crosstalk, was also present within the MYC2 subnetwork (Supplementary Fig. 6a), predicted to inhibit MYC2 but not to be reciprocally regulated by MYC2 (Wild et al., 2012). These three interactions are potential points at which crosstalk can occur rapidly during a JA response with auxin, gibberellin and ethylene.

We next examined the broader gene regulatory network to identify additional predicted points of crosstalk between JA and other signaling pathways. The model predicted that STZ/ZAT10 is a key early hub through which JA signaling is prioritized over several other hormone and stress response pathways (Fig. 5a and Supplementary Table 12). STZ/ZAT10 is known to be a transcriptional repressor from genetic studies (Mittler et al., 2006) and, consistent with this, our model predicted that it inhibited the majority of genes it regulates (25 of 34 genes). *WRKY40*, *WRKY70*, *DDF* and *ERF6* were all predicted to be inhibited by STZ/ZAT10 within 0.25 h of a JA stimulus and *GRX480* within 1 h. Direct binding of STZ/ZAT10 to ERF6 was detected in ChIP-seq assays (Supplementary Table 6). WRKY40 and WRKY70 both are both early brassinosteroid response components that repress defense responses (Lozano-Duran et al., 2013). WRKY70 also fine-tunes the crosstalk between the salicylic acid and JA pathways (Li et al., 2006). DDF1 promotes resistance to drought, cold, heat and salinity stress by reducing endogenous gibberellin abundance (Kang et al., 2011, Magome et al., 2008). ERF6 similarly promotes drought resistance by reducing gibberellin abundance (Dubois et al., 2015). GRX480 regulates the negative crosstalk between salicylic acid and both JA/ethylene signaling through the direct interactions with TGA transcription factors (Zander et al., 2012, Ndamukong et al., 2007). The model also predicts that ERF6, WRKY70 and DDF1 exert negative feedback on STZ/ZAT10 by activating JAZ8 within 0.25 h of the JA stimulus (Fig. 5a and Supplementary Table 12). JAZ8 is a known repressor of JA signaling and is predicted to repress STZ/ZAT10 (Shyu et al., 2012). In sum the gene regulatory network predicts that STZ/ZAT10 is an important hub for JA signaling to be prioritized over other hormone and stress response pathways (Fig. 5a).

**Figure 5.**
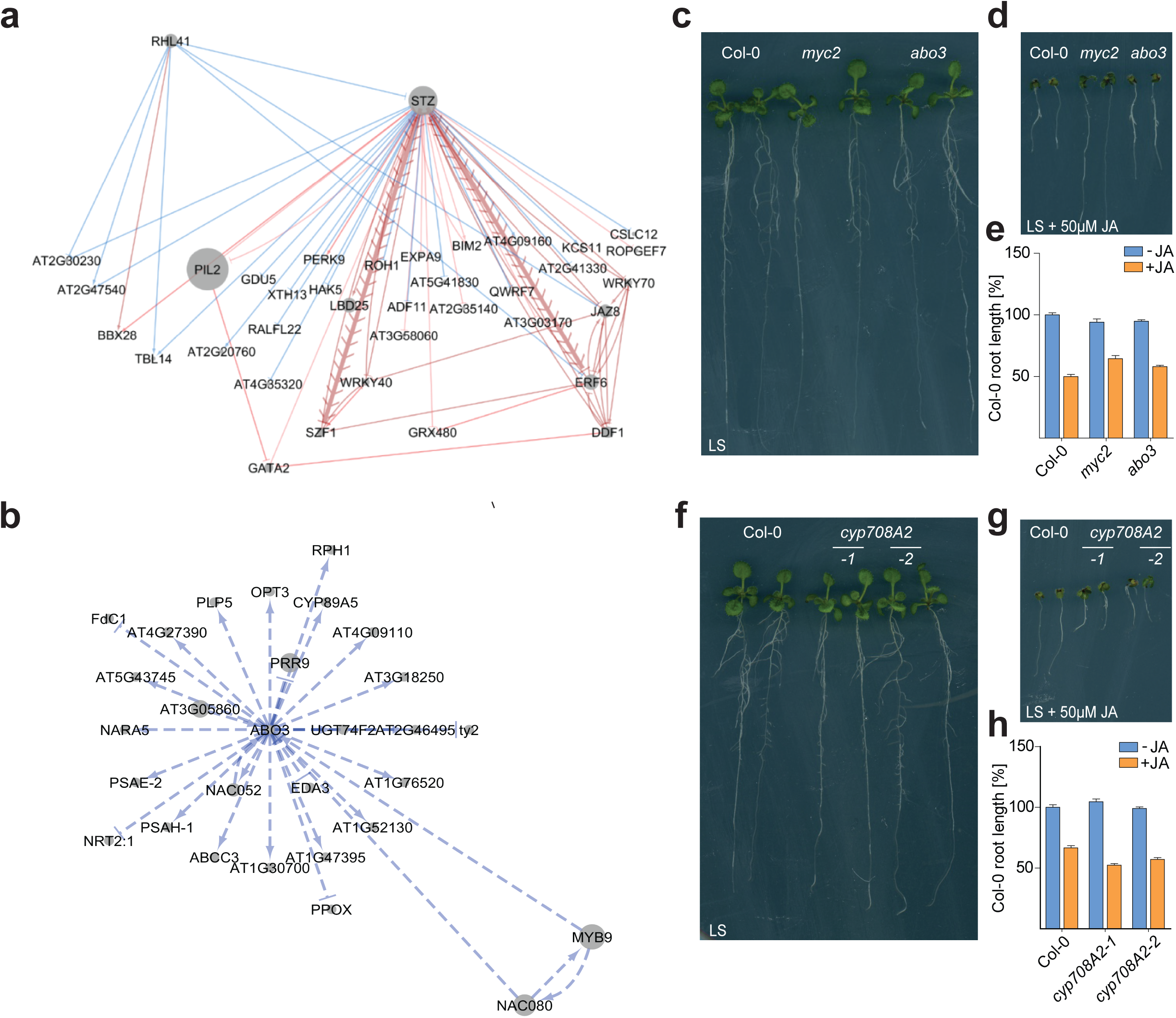
JA response genome regulatory model positions known and new components. **a, b,**Subnetworks of STZ (a) and ABO3 (b) are shown. Edges are directed. Red edges exist at early time points (0.25 – 2 h), blue only at late time points (4 – 24 h). Thicker edges with chevrons indicate a MYC2 directly bound that gene in ChIP-seq experiments. **c, d,** JA-induced root growth inhibition assay identified ABO3 as a positive JA regulator. Seedlings were grown on LS media with (d) or without 50µM MeJA (c). Col-0 and *myc2* seedlings served as controls. **e,** Quantification of JA-induced root growth inhibition in Col-0 and *abo3* seedlings is shown. **f, g,** Root growth inhibition assay identified two *cyp708A2* T-DNA alleles as JA hyper-sensitive. Seedlings were grown on LS media with (f) or without 50µM MeJA (g) and Col-0 and *myc2* seedlings serve as controls. **h,** Bar plot shows quantification of JA-induced root growth inhibition in Col-0 and *cyp708A2* seedlings.

### Phenotypic screening guided by large-scale data identifies new JA signaling components and validates the JA gene regulatory network

We next utilized our regulatory network and large-scale datasets to identify novel regulators of the JA pathway using the JA root growth inhibition assay as our experimental readout. First, we focused on ABO3 (ABA overly sensitive 3), which is directly targeted by MYC2 and MYC3 (Fig. 2d and Supplementary Table 1) and whose subnetwork is comprised of 26 predicted regulated genes, the majority of which is positively regulated (22 of 26 genes) (Fig. 5b). *ABO3* encodes the *Arabidopsis* WRKY transcription factor gene WRKY63, which is involved in stress gene expression and drought tolerance (Ren et al., 2010, Van Aken et al., 2013). To investigate the importance of the ABO3 subnetwork in JA signaling, we tested *abo3* T-DNA mutant seedlings (SALK_075986C) in a JA-induced root growth inhibition assay. We found that *abo3* mutants show a weak JA hypo-sensitive root growth inhibition phenotype (Fig. 5c-e) indicating that ABO3 is positive regulator of JA signaling and that our network approach is able to identify new pathway components.

Next, we expanded our phenotyping analysis to T-DNA lines of genes that display the strongest binding of MYC2 and MYC3 in their promoters (Supplementary Table 1, 13). The rationale behind this approach is that master TFs target the majority of key signaling components in their regulated respective pathways and that these are often the most strongly bound targets (Chang et al., 2013, Song et al., 2016, Xie et al., 2018). Of 99 genes tested (194 T-DNA lines in total, Supplementary Table 14), we discovered six genes, when mutated, display mild JA root growth phenotypes (Supplementary Fig. 7a and Supplementary Table 14). Mild phenotypes as well as their low frequency were not surprising since gene redundancy is very common in the *Arabidopsis* genome and even the mutation of the master TF MYC2 only causes a mild JA-hyposensitive root growth phenotype (Fig. 5c-e) (Lorenzo et al., 2004, 2000). Among these genes was the cytochrome P450 enzyme *CYP708A2* gene from which both tested T-DNA mutant alleles exhibit a JA hypersensitive root phenotype (Fig. 5f-h). Interestingly, our network analysis also discovered CYP708A2 as a regulatory hub (Supplementary Fig. 5a, 7b). CYP708A2 is involved in the triterpene synthesis which is known to be stimulated by methyl jasmonate (Field and Osbourn, 2008, Mangas et al., 2006); future studies are however needed to further decipher the role of CYP708A2 in JA signaling. Another interesting uncharacterized gene that we discovered caused a JA phenotype is a Sec14p-like phosphatidylinositol transfer family protein (*AT5G47730*) (Supplementary Fig. 7a and Supplementary Table 14). Phosphatidylinositol transfer proteins (PITPs) are crucial for the phosphatidylinositol homeostasis in plants (Huang et al., 2016) and inositol polyphosphates have been implicated in COI1-mediated JA perception (Mosblech et al., 2011). Taken together, these data show that our multi-omic approach goes beyond network description ultimately enabling the identification of novel JA pathway regulators.

## Discussion

An important unanswered question in plant biology is how multiple signaling pathways interact to coordinate control of growth and development. In this study we have comprehensively characterized the cellular response to the plant hormone JA and generated a network-level understanding of the MYC2/MYC3-regulated JA signaling pathway. We used this to identify several new points at which JA signaling may have cross-regulation with other hormone and stress response pathways in order to prioritize itself. The results increase knowledge of how JA functions in the etiolated seedling, a less well characterized model for JA responses. Moreover, the general principles described here provide a framework for analysis of cross-regulation between hormone and stress signaling pathways. We provide our data in a web-based genome and network browsers to encourage deeper exploration (http://signal.salk.edu/interactome/JA.php, http://neomorph.salk.edu/MYC2).

The major insight provided by our study is that multiple points of crosstalk are likely to exist between JA signaling and other pathways. This was evident from the interactions within the genome regulatory network model and supported by our observation that many (37 to 59%) genes from other hormone signaling pathways are bound by MYC2/3 and JA-regulated. The WRKY family TF ABO3 was identified as a candidate JA response regulator and genetic analyses determined a mutant of the gene was JA hyposensitive. ABO3 is also a regulator of ABA responses (Ren et al., 2010) suggesting that ABO3 functions in the cross-communication between the JA and ABA pathway. The repressive zinc-finger family TF STZ/ZAT10, working with JAZ8, emerged as a potentially important point of contact with salt and drought stress, as well as the salicylic acid, brassinosteroid and gibberellin hormone signaling pathways. Combined these results illustrate the importance of transcriptional cross-regulation during a JA response in modulating the correct cellular output for the stimuli a plant perceives.

Our multi-omic analysis determined that the master TF MYC2 and its relative MYC3 directly target thousands of JA responsive genes including hundreds of JA responsive TFs, thereby enabling a robust cascade of transcriptional reprogramming. Secondary TFs downstream of MYC2/3 directly targeted overlapping but distinct cohorts of genes, indicating they have distinct roles within the JA response. This illustrates the complexity of hormone-response genome regulatory programs; we have assayed only a fraction of the JA-responsive TFs and find that any individual JA-responsive gene may be bound by multiple TFs. How the final quantitative output of any individual gene is determined by combinatorial binding of TFs remains a major challenge to address. We further demonstrated the importance of MYC2/3 target genes in JA responses by analyzing JA root growth phenotypes in mutants of 99 genes strongly targeted by MYC2/3. Mutations in seven genes caused clear disruption of JA responses, both hyper and hypo-sensitivity. It is probable that genetic redundancy accounts for a proportion of the mutants not causing phenotype changes.

Another layer of regulatory complexity within the JA signaling pathway, and within signaling pathways in general, is the presence of multiple feedforward and feedback loops that are activated simultaneously. The interactions between these subnetworks through their kinetics and the strength of their regulatory impact on the broader network is not well understood. For example, we discovered that MYC2 and MYC3 stimulate JA biosynthesis but also target the entire JAZ repressor family from which the majority of members is also transcriptionally activated. Uncoupling these subnetworks would be an effective way to determine how they interact to drive very robust activation of the JA pathway. The combination of our multi-omic framework approach coupled with powerful genetic approaches such as the generation of the *jaz* decuple mutant (Guo et al., 2018) should significantly contribute to a better understanding of JA response pathways

## Supporting information

All supplemental figures

## Acknowledgements

We thank several postdocs, undergrads and technicians who contributed technical assistance to the project; Mingtang Xie, Liang Song, Raul Carlos Serrano, Candice Sy, Lourdes Tames, Julie Park, Omar Romero, Raymond Luong, Waina Ho, Yusuke Koga, Sasha Hazelton, Mark Urich, Tsegaye Dabi. We thank Shao-shan Carol Huang for computational assistance and James Moresco and Jolene Diedrich for proteomics support.

## Contributions

M.Z., M.G.L., R.S. and J.R.E. designed the research. M.Z., M.G.L., A.E.L. and B.J. performed the phenotype screening. M.Z., M.G.L. and J.P.S.G. carried out the RNA-seq and ChIP-seq experiments. M.G.L., E.H. and J.P.S.G. performed the cloning and generation of transgenic constructs. M.G.L., J.R.N., H.C, M.Z. and L.Y. analyzed the sequencing data and performed bioinformatics analyses. A.B. carried out DAP-seq experiments. N.M.C. and J.W.W. analyzed the proteome and phosphoproteome data. N.M.C., J.W.W., A.W., S.J. and Z. B-J. performed regulatory network analyses. M.Z., M.G.L and J.R.E. prepared the figures and wrote the manuscript.

## Data and material availability

All described lines can be requested from the corresponding author. Sequence data can be downloaded from GEO (GSE133408). Proteomics data are deposited at Proteome Exchange under the accession ID PXD013592. Visualized data can be found under http://neomorph.salk.edu/MYC2 and http://signal.salk.edu/interactome/JA.php.

## Material & Methods

### Plant material and growth conditions

The *myc2* mutant in this study is *jin1-8* (SALK_061267) (Lorenzo et al., 2004) and was obtained from the Arabidopsis Biological Resource Center (ABRC). Col-0 *MYC2::MYC2-Ypet* and Col-0 *MYC3::MYC3-Ypet*, generated by recombineering, have been described previously (Zhou et al., 2011). For the generation of all large scale datasets, three-day-old etiolated seedlings were used (Col-0, *myc2*, *MYC2::MYC2-Ypet*, *MYC3::MYC3-Ypet*). Gaseous MeJA treatment for the respective times was performed as previously described (Schweizer et al., 2013b). For the JA-induced root growth inhibition assay, surface-sterilized Col-0, *myc2* and T-DNA mutant seeds (Supplementary Table 13) were grown on agar plates containing LS medium supplemented with or without 50 µM MeJA (392707, Millipore Sigma) for 9 days. Plates were scanned afterwards and root length was measured using ImageJ.

### ChIP-seq

Three-day-old etiolated Col-0 *MYC2::MYC2-Ypet*, Col-0 *MYC3::MYC3-Ypet,* Col-0 and *myc2* seedlings were used for ChIP-seq experiments. ChIP assays were performed as previously described (Kaufmann et al., 2010). ChIP-seq assays were conducted with antibodies against H2A.Z (39647, Active Motif), H3K4me3 (04-745, Millipore Sigma) and GFP (11814460001, Millipore Sigma or goat anti-GFP supplied by David Dreschel, Max Planck Institute of Molecular Cell Biology and Genetics). As a negative control, mouse or goat IgG (015-000-003 or 005-000-003, Jackson ImmunoResearch) was used. The respective antibodies or IgG were coupled for 4-6 hour to Protein G Dynabeads (50µl, 10004D, Thermo Fisher Scientific) and subsequently incubated overnight with equal amounts of sonicated chromatin. Beads were washed twice with high salt buffer (50 mM Tris HCl pH 7.4, 150 mM NaCl, 2 mM EDTA, 0.5% Triton X-100) low salt (50 mM Tris HCl pH 7.4, 500 mM NaCl, 2 mM EDTA, 0.5% Triton X-100) and wash buffer (50 mM Tris HCl pH 7.4, 50 mM NaCl, 2 mM EDTA) before samples were de-crosslinked, digested with proteinase K and DNA was precipitated. Sequencing libraries were generated following the manufacturer’s instructions (Illumina). Libraries were sequenced on the Illumina HiSeq 2500 and HiSeq 4000 Sequencing system and sequencing reads were aligned to the TAIR10 genome assembly using Bowtie2 (Langmead, 2010).

### DAP-seq

DAP-seq assays were carried as previously described (O’malley et al., 2016) using recombinantly expressed ERF1 (AT3G23240, ERF1B, AtERF092), ORA59 (AT1G06160), ATAF1 (AT1G01720), DREB2B (AT3G11020), ZAT18 (AT3G53600), RVE2 (AT5G37260), WRKY51 (AT5G64810), HY5 (AT5G11260) and TCP23 (AT1G35560).

### RNA-seq

Three-day-old etiolated seedlings were used for expression analyses. Total RNA was extracted with the RNeasy Plant Mini Kit (74903, Qiagen). cDNA library preparation and subsequent single read sequencing was carried as previously described (Song et al., 2016).

### RNA-seq analyses

Sequencing reads were quality trimmed using TrimGalore 0.4.5 https://github.com/FelixKrueger/TrimGalore) then aligned to the TAIR10 genome assembly using TopHat 2.1.1 (Kim et al., 2013). Reads within gene models were counted using HTSeq (Anders et al., 2015). Differentially expressed genes in time series RNA-seq were identified using EdgeR 3.6.2 with a likelihood ratio test (functions glmFit and glmLRT), batch correction Benjamini & Hochberg correction for multiple tests (Robinson et al., 2010). Differentially expressed genes in the Col-0 *versus myc2* mutant RNA-seq were determined using EdgeR 3.18.1 and quasi-likelihood F-tests (function glmQLFit) (Lun et al., 2016). Temporal co-regulation of transcripts was determined using the Short Time-Series Expression Miner (Ernst and Bar-Joseph, 2006). Known *A. thaliana* TFs were identified by reference to PlantTFDB 4.0 (Jin et al., 2017).

### ChIP-seq and DAP-seq analyses

ChIP-seq and DAP-seq sequence reads were mapped to the TAIR10 reference genome using Bowtie 2 v.2-2.0.5 with default parameters (Langmead and Salzberg, 2012). For TF ChIP-seq, enriched binding sites were identified using MACS2 v.2.1 (options -p 99e-2 --nomodel –shiftsize--down-sample --call-summits) against sequence reads from whole IgG control samples (Zhang et al., 2008). The shift size was calculated using PhantomPeakQualTools v.2.0 (Kharchenko et al., 2008). Subsequent analyses used summits only. Summit lists were filtered with a lower cut-off of -log10(25) and remaining summits expanded from single nucleotides to 150 nt. Only summits with at least 10% nt overlap between at least two biological replicates were retained. These overlapping summits were merged between replicates using BEDtools v.2.17.0 to give the final set of high-stringency summits, which were then annotated using ChIPpeakAnno v.2.2.0 to any gene within 500 nt of the center of the summit or, alternatively, the nearest neighbor if there was no gene within 500 nt (Quinlan and Hall, 2010, Zhu et al., 2010). Venn diagrams were drawn using Venny and Intervene (http://bioinfogp.cnb.csic.es/tools/venny/) (Khan and Mathelier, 2017). Top-ranked MYC2/3 binding sites were identified by applying IDR to the summits from the two biological replicates that had the greatest number of summits above the MACS2 lower cut-off of -log10(25). TF binding motifs were determined using the MEME-ChIP webserver with default parameters on the sequences of the high-stringency summits (Machanick and Bailey, 2011). The Genome wide Event finding and Motif discovery (GEM) tool (Guo et al., 2012) was used to identify the target summits in DAP-seq data. Significant enrichments of histone modifications and histone variants were identified with the SICER software (Zang et al., 2009) using the TAIR10 genome assembly. The Intersect tool from BEDtools (Quinlan and Hall, 2010) was used to identify the genes in the ChIP-seq datasets that are most proximal to the discovered binding sites. For both ChIP-seq and DAP-seq gene ontology enrichment was assessed using clusterProfiler (Yu et al., 2012).

### Mass spectrometry analysis

Ground untreated/JA-treated Col-0 and *myc2* seedlings tissue was ground and lysed in YeastBuster (71186, Millipore Sigma). Proteins (100 µg per sample) were precipitated using methanol-chloroform. Dried pellets were dissolved in 8 M urea, 100 mM triethylammonium bicarbonate (TEAB), reduced with 5 mM tris (2-carboxyethyl) phosphine hydrochloride (TCEP), and alkylated with 50 mM chloroacetamide. Proteins were then trypsin digested overnight at 37°C. The digested peptides were labeled with TMT10plex™ Isobaric Label Reagent Set (90309, Thermo Fisher Scientific, lot TE264412) and combined. One hundred micrograms (the pre-enriched sample) was fractionated by basic reverse phase (84868, Thermo Fisher Scientific). Phospho-peptides were enriched from the remaining sample (900 µg) using High-Select Fe-NTA Phospho-peptide Enrichment Kit (A32992, Thermo Fisher Scientific). The TMT labeled samples were analyzed on a Fusion Lumos mass spectrometer (Thermo Fisher Scientific). Samples were injected directly onto a 25 cm, 100 μm ID column packed with BEH 1.7 μm C18 resin (186002350, Waters) and subsequently separated at a flow rate of 300 nL/min on a nLC 1200 (LC140, Thermo Fisher Scientific). Buffer A and B were 0.1% formic acid in water and 90% acetonitrile, respectively. A gradient of 1–20% B over 180 min, an increase to 40% B over 30 min, an increase to 100% B over another 20 min and held at 90% B for a final 10 min of washing was used for 240 min total run time. Column was re-equilibrated with 20 μL of buffer A prior to the injection of sample. Peptides were eluted directly from the tip of the column and Nano sprayed directly into the mass spectrometer by application of 2.8 kV voltage at the back of the column. The Lumos was operated in a data dependent mode. Full MS1 scans were collected in the Orbitrap at 120000 resolution. The cycle time was set to 3 s, and within this 3 s the most abundant ions per scan were selected for CID MS/MS in the ion trap. MS3 analysis with multinotch isolation (SPS3) was utilized for detection of TMT reporter ions at 60000 resolution. Monoisotopic precursor selection was enabled and dynamic exclusion was used with exclusion duration of 10 s.

The raw data were analyzed using MaxQuant version 1.6.3.3 (Tyanova et al., 2016). Spectra were searched, using the Andromeda search engine (Cox et al., 2011) against the Tair10 proteome file entitled “TAIR10_pep_20101214” that was downloaded from the TAIR website (https://www.arabidopsis.org/download/indexauto.jsp?dir=%2Fdownload_files%2FProteins%2FTAIR10_protein_lists) and was complemented with reverse decoy sequences and common contaminants by MaxQuant. Carbamidomethyl cysteine was set as a fixed modification while methionine oxidation and protein N-terminal acetylation were set as variable modifications. For the phoshoproteome “Phosho STY” was also set as a variable modification. The sample type was set to “Reporter Ion MS3” with “10plex TMT selected for both lysine and N-termini”. Digestion parameters were set to “specific” and “Trypsin/P;LysC”. Up to two missed cleavages were allowed. A false discovery rate, calculated in MaxQuant using a target-decoy strategy (Elias and Gygi, 2010) less than 0.01 at both the peptide spectral match and protein identification level was required. The ‘second peptide’ option identify co-fragmented peptides was not used. Differentially expressed proteins and phospho-sites were identified using PoissonSeq (Li et al, 2012) with a q-value cutoff of 0.1. Sample loading normalization was performed before differential expression analysis.

### Transcript quantification and identification of isoform switches

Quantification of transcripts was performed using Salmon v0.8.1 in conjunction with the AtRTD2-QUASI transcript reference (Patro et al., 2017, Zhang et al., 2015b) The quasi mapping-based index was built using an auxiliary k-mer hash over k-mers of length 31 (k=31). For quantification, all parameters of Salmon were kept at default, except that the option to correct for the fragment-level GC biases (“–gcBias”) was turned on.

The TSIS R package, which is designed for detecting alternatively spliced isoform switch events in time-series transcriptome data, was used to perform the isoform switch analysis (Guo et al., 2017) Only transcripts whose average transcript per million (TPM) across all time points was >1 were included in the TSIS analysis. The mean expression approach was used to search interaction points. Significant switch events were identified using the following filtering parameters: (1) probability cutoff >0.5; (2) differences cutoff >1; (3) p-value cutoff < 0.05; (4) min

### Gene regulatory network (GRN) inference

All GRNs were constructed using the Regression Tree Pipeline for Spatial, Temporal, and Replicate data (RTP-STAR) (Shibata et al., 2018) Clark et al., 2018). Prior to GRN inference, genes were clustered based on transcriptome, proteome, or phosphoproteome data using Dynamic Time Warping (DTW) (and the dtwclust package in R (Giorgino 2009). These clusters were then used in the RTP-STAR pipeline. For the transcriptome networks, one network was inferred for genes differentially expressed at each time point (8 networks total), and then the networks were combined in a union. For each network, the biological replicates for that individual time point and the 0 h (control) time point were used to infer the network. The sign (activation/repression) of each edge was inferred using all of the time points in the time course. For the proteome and phosphoproteome networks, one network was inferred for genes differentially expressed in any of the comparisons. The biological replicates for all of the (phospho) proteome samples were used to infer the network. The sign of each edge was not inferred as the (phospho) proteome data only consisted of one time point.

After the transcriptome, proteome, and phosphoproteome networks were combined in a union, a Network Motif Score (NMS; Clark et al., 2018) was calculated to determine the importance of each gene. Feedback loop, feed-forward loop, bi-fan, and diamond motifs were used in this score as they have been previously shown to contain genes important for biological processes (Alon, 2007, Milo et al., 2002, Ingram et al., 2006). All motifs were significantly enriched in the combined network compared to a randomly generated network of the same size. The number of times each gene appeared in each motif was counted, the counts were normalized to a scale of 0 to 1, and the counts were summed to calculate the NMS. The higher the NMS, the more functionally important the gene is. All code for RTP-STAR is available at https://github.com/nmclark2/RTP-STAR. The parameters used for all networks in this paper are provided in Supplementary Table 15.

## Supplementary Figures

**Supplementary Figure 1. MYC2 and MYC3 regulate the majority of JA signaling pathway components**

**a,** Gene ontology (GO) analysis of MYC2 and MYC3 targets is shown. Analysis was conducted using clusterProfiler. **b,** Bar plots shows the portion of JA-induced and JA-repressed genes that are bound by MYC2 (b) and MYC3 (c). **c,** Binding behavior of MYC2 and MYC3 at known JA genes (Supplementary Table 4) is shown. Known JA genes are grouped into non-differentially expressed and JA differentially expressed genes. **d,** Schematic overview of known MYC2/MYC3-targeted JA pathway components. **e,** AnnoJ genome browser screenshot visualizes MYC2 and MYC3 binding at all 13 members of the JAZ repressor family, as well as at the co-repressor adaptor protein NINJA. mRNA expression of these genes in untreated/JA-treated Col-0 and *myc2* seedlings is also shown.

**Supplementary Figure 2. MYC2 and MYC3 target a large number of TFs**

**a.** Cluster analysis revealed the 5 other main clusters in the JA time course experiment. Clusters visualize the log2 fold change expression dynamics over the indicated 24 hours’ time period. The three strongest enriched gene ontology terms for each cluster are shown as well. **b,** Pie Chart indicates the proportions of TFs that are transcriptionally induced by JA, bound by MYC2/MYC3, or both. **c, d,** Overview of MYC2/MYC3-bound plant hormone genes (c) and TFs (d) is shown. Plant hormones are abbreviated (ET (ethylene), BR (brassinosteroids), GA (gibberellic acid), ABA (abscisic acid), SA (salicylic acid), CK (cytokinin), AUX (Auxin), K (karrikin), SL (strigolactones)).

**Supplementary Figure 3. Overview of MYC-controlled TF network**

Significantly enriched (adjusted p<0.05) gene ontology terms amongst the target of each TF. For each TF the 4 terms with the lowest p-value are shown, some of which are redundant between TFs. No enriched terms were detected for DREB2B targets.

**Supplementary Figure 4. Jasmonic acid shapes the local chromatin architecture**

**a,** Bar plot shows the impact of two hours JA treatment on the genome-wide distribution of H3K4me3 and H2A.Z domains. Occupancy was determined in untreated/JA-treated Col-0 and *myc2* seedlings using ChIP-seq. SICER was used to identify the number of histone domains that show an increase (blue) or decrease (orange) of enrichment in response to JA. **b, c,** Heatmaps show the occupancy of H3K4me3 and H2A.Z from 1 kb upstream to 2 kb downstream of the transcriptional start site (TSS) at all *Arabidopsis* genes (TAIR10). Heatmaps are shown for H3K4me3 (b) and H2A.Z (c) in untreated and JA-treated (4 h) Col-0 and *myc2* seedlings. **d,** Quantification of H3K4me3 and H2A.Z occupancy at *JAZ2* and *GRX480* are shown. It was calculated as the ratio between the respective ChIP-seq sample and the Col-0 IgG control.

**Supplementary Figure 5. The jasmonic acid gene regulatory network**

**a,** Illustration of JA gene regulatory network for 1, 2 and 4 h time points. Edges were predicted using phosphoproteome (green), proteome (orange) and transcriptome (blue) data. Node sizes are scaled by normalized motif score, with larger nodes indicating greater scores and likely greater importance within the network. Edges predicted early in the time-series transcriptomic data are red (0.25 – 2 h), edges predicted late are blue (4 - 24 h). Proteome and phosphoproteome-data-predicted edges are grey and green, respectively.

**Supplementary Figure 6. Gene regulatory network validation against ChIP/DAP-seq data a,** The MYC2 subnetwork is shown. Edges are directional and red edges exist at early time points (0.25 – 2 h), blue only at late time points (4 – 24 h). Thicker edges with chevrons indicate that MYC2 were directly bound to that gene in our ChIP-seq experiments. **b,** Validated edges are those between TFs and first neighbours in the JA gene regulatory network for which the first neighbour was also a direct target of the TF in ChIP/DAP-seq assays. These edges are indicated by chevrons. Early time-series transcriptome-predicted edges (0.25 – 2 h) are red and later edges (4 - 24 h) are blue. Edges detected in the proteomic data are grey and those detected in the phosphoproteomic data are green.

**Supplementary Figure 7 Validation of regulatory network predictions**

**a,** Bar plot shows quantification of JA-induced root growth inhibition in the indicated T-DNA alleles. Seedlings were grown on LS media with or without 50µM MeJA. Col-0 seedlings serve as independent controls for each tested T-DNA line. **b,** Subnetwork of CYP708A2 is shown.

## Supplementary Tables

**Supplementary Table 1.** High-confidence target genes of MYC2 and MYC3 after two hours JA treatment. Unprocessed summit outputs of biological replicate experiments including p-value scores. Hormone transcription factors bound by MYC2/3.

**Supplementary Table 2.** Differential regulation of all transcripts relative to 0 h abundance following JA treatment. Tab names indicate time point post-treatment. Calculated by EdgeR with false discovery rate (FDR)<0.05 indicating statistical significance. FC - fold change. CPM - counts per million. LR - likelihood ratio.

**Supplementary Table 3.** Motifs detected *de novo* within MYC2 and MYC3 target summits. The data are DREME model outputs for MYC2 and MYC3 high-stringency summits.

**Supplementary Table 4.** Expression of 168 known JA genes following JA stimulus and whether they are bound by MYC2/3 or not. ND indicates non-differentially regulated, as assessed by EdgeR (FDR<0.05).

**Supplementary Table 5.** Details of STEM model of JA-responsive transcripts and details of transcripts within statistically significant clusters. Input data were the expression values of all transcripts significantly differentially regulated at any time in the time series relative to 0 h post-JA stimulus.

**Supplementary Table 6.** Target genes of JA transcription factors identified by ChIP-seq (NAC3/ANAC055, STZ/ZAT10; indicated by *) and DAP-seq (DREB2B, ATAF2, HY5, RVE2, ZAT18, TCP23, ERF1, ORA59, WRKY51).

**Supplementary Table 7.** Differentially enriched H3K4me3 and H2A.Z domains in JA-treated Col-0 and *myc2* seedlings.

**Supplementary Table 8.** Differentially expressed proteins detected in proteomics analyses.

**Supplementary Table 9.** Differentially abundant phosphopeptides detected in phosphoproteomic analyses.

**Supplementary Table 10.** Transcript pairs exhibiting isoform switch events as detected by TSIS analyses.

**Supplementary Table 11.** TPM quantification of transcripts in the JA time-series RNA-seq against the AtRD2 reference transcriptome.

**Supplementary Table 12.** Nodes and edges within the JA response genome regulatory network model, generated from combined JA (phospho)proteome and transcriptome data. Normalized motif score of all components is also given.

**Supplementary Table 13.** Gene regulatory network validation against ChIP/DAP-seq data. Validated edges are those between TFs and first neighbours in the JA gene regulatory network for which the first neighbour was also a direct target of the TF in ChIP/DAP-seq assays. * indicates ChIP-seq assay, all others were DAP-seq.

**Supplementary Table 14.** List of tested T-DNA lines and T-DNA lines with a JA root growth inhibition phenotypes.

**Supplementary Table 15.** List of parameters used during gene regulatory network reconstruction.

