## Supplementary material for "Integrated Multi-omic Framework of the Plant Response to Jasmonic Acid": All supplemental figures

### Supplementary Figure 1

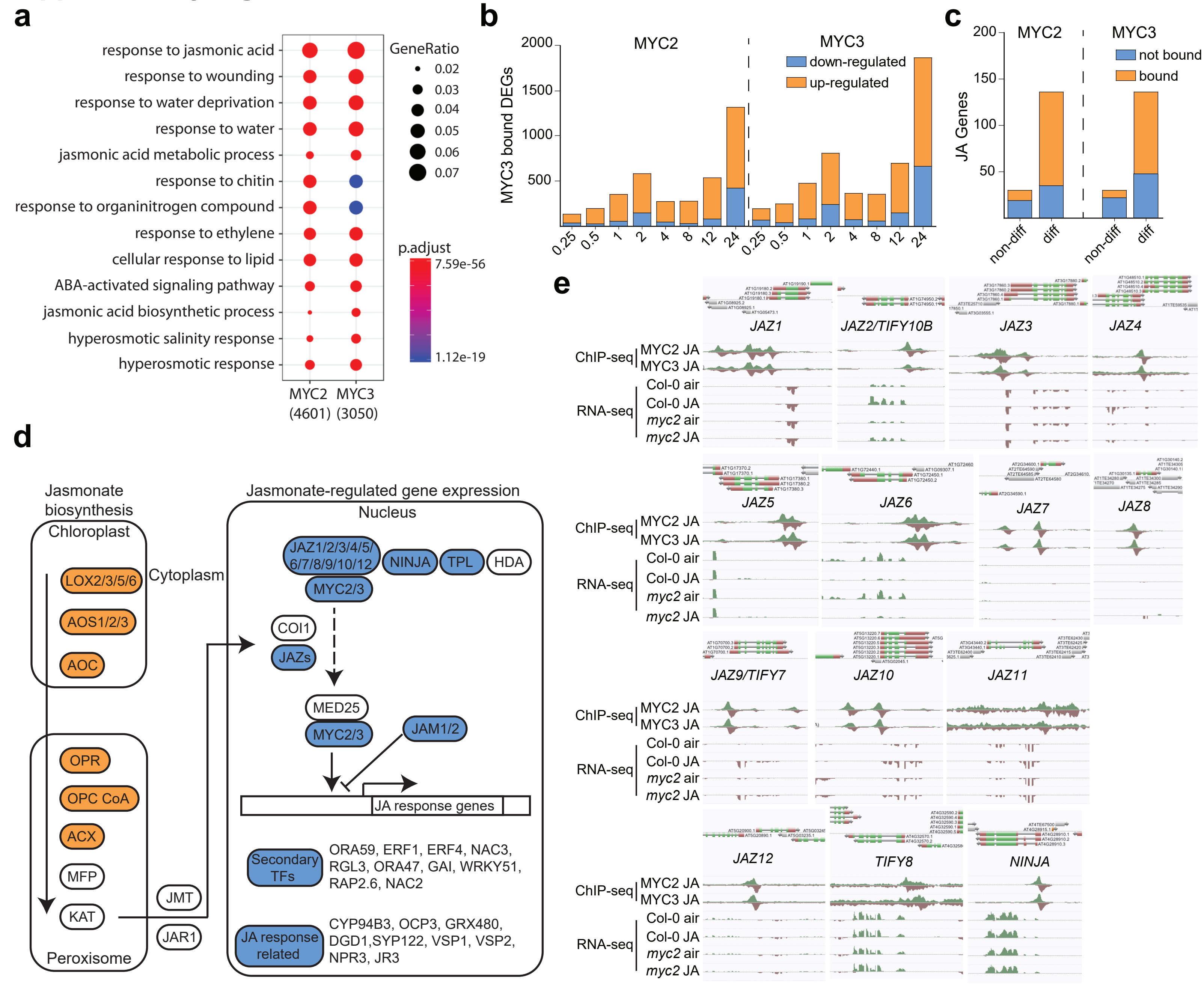

### Supplementary Figure 2

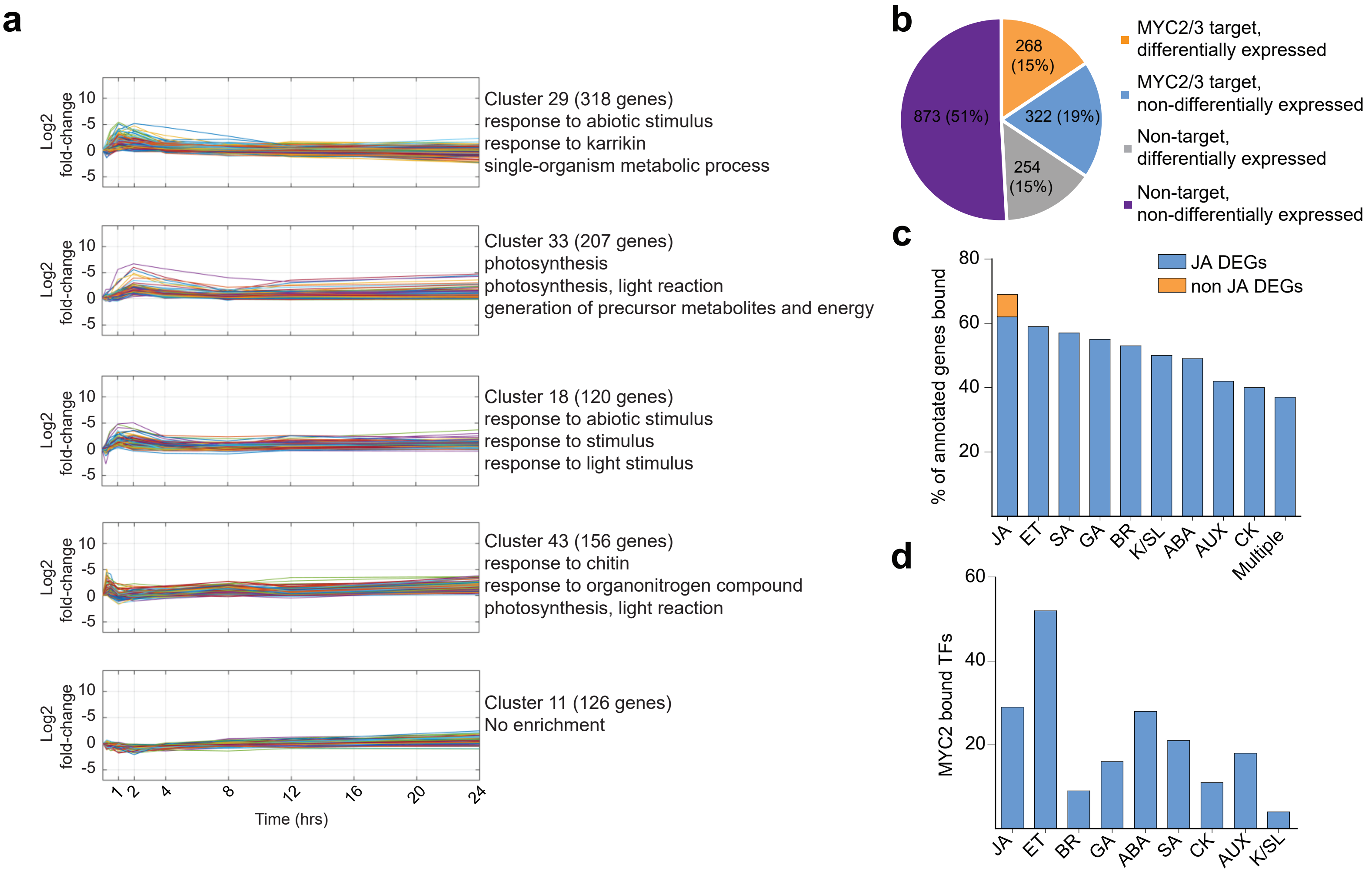

### Supplementary Figure 3

## a

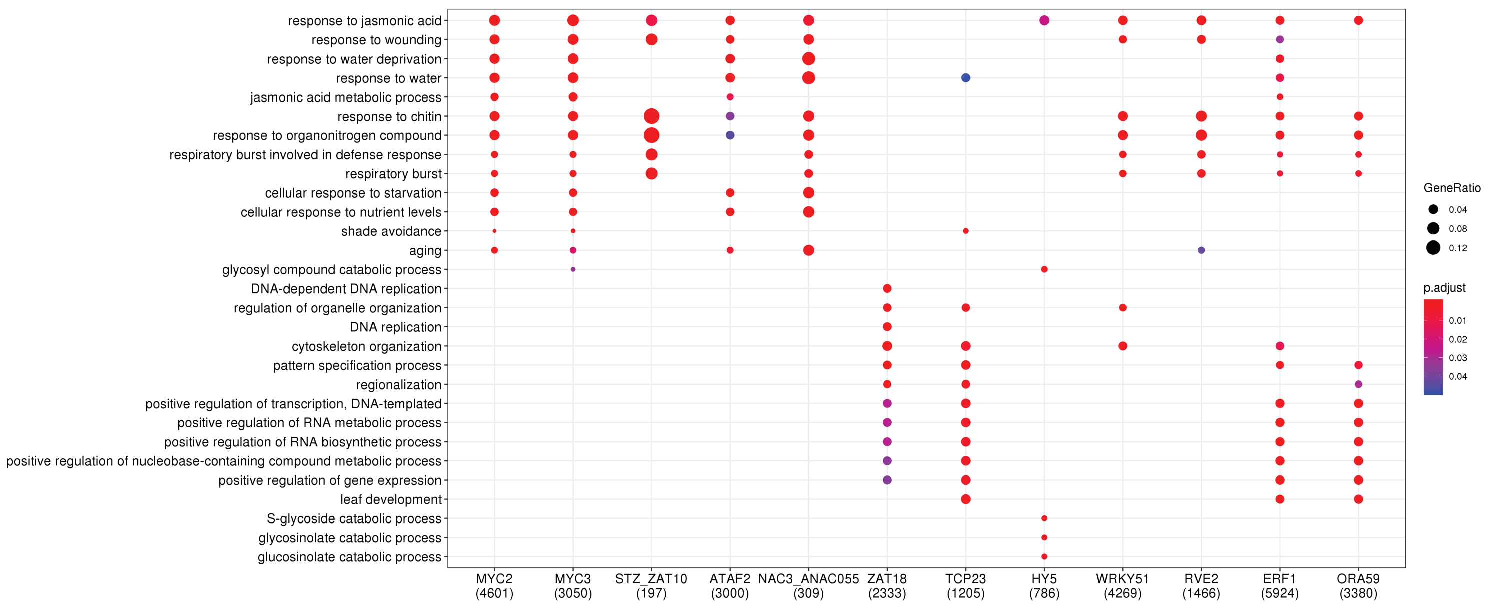

### Supplementary Figure 4

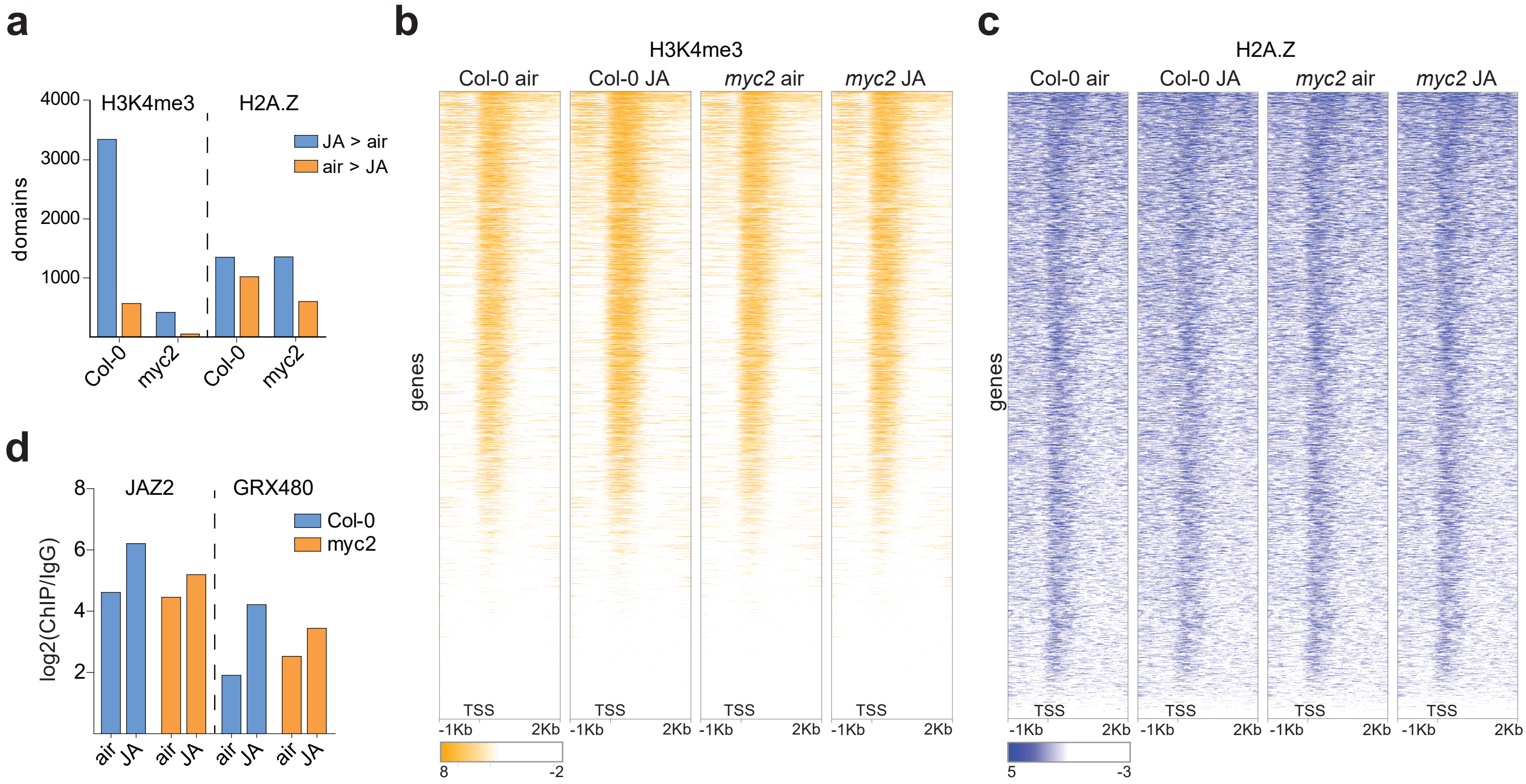

#### Supplementary Figure 5

**a**

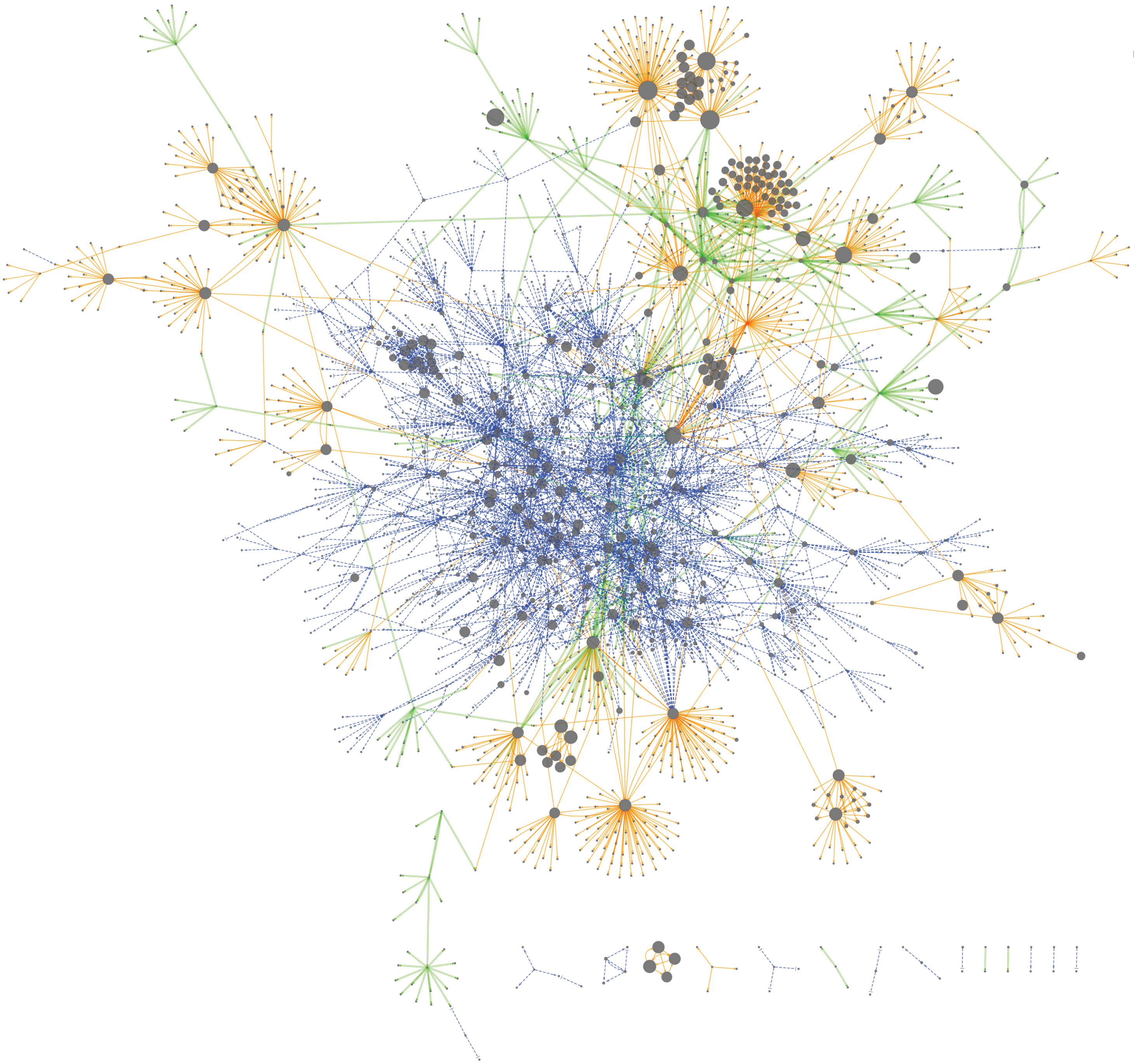

### Supplementary Figure 6

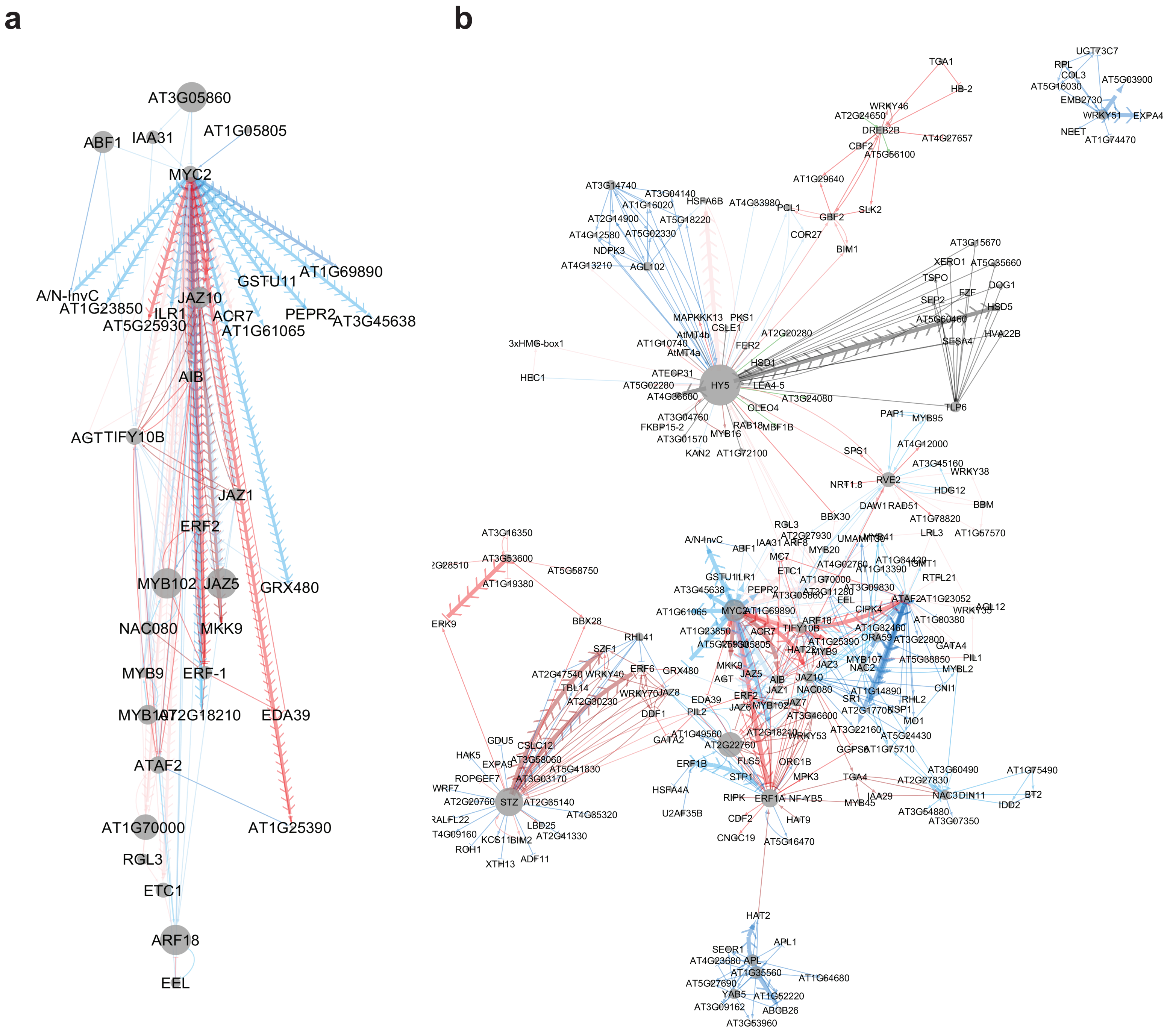

### Supplementary Figure 7

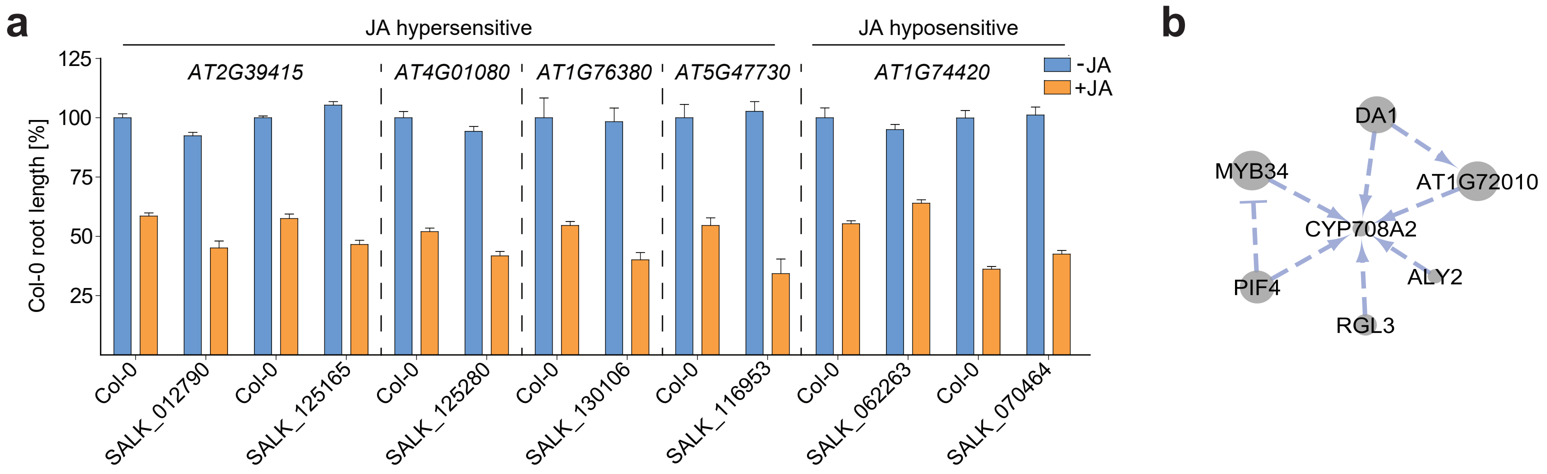
